# Th17 cells provide direct protective effects that limit stomach parasite burden following orogastric mucosal *Trypanosoma cruzi* infection

**DOI:** 10.1101/2020.11.24.397257

**Authors:** Catherine W. Cai, Christopher S. Eickhoff, Krystal A. Meza, Jennifer R. Blase, Rebecca E. Audette, David H. Chan, Kevin A. Bockerstett, Richard J. DiPaolo, Daniel F. Hoft

## Abstract

*Trypanosoma cruzi* is the intracellular parasite of Chagas disease, a chronic condition characterized by cardiac and gastrointestinal morbidity. Protective immunity requires CD4+ T cells, and Th1 cells and IFN-γ are important players in host defense. More recently, Th17 cells and IL-17 have been shown to exert protective functions in systemic *T. cruzi* infection. However, it remains unclear whether Th17 cells and IL-17A protect against mucosal infection, which is an important cause of human outbreaks. We found that IL-17RA knock-out (KO) mice are highly susceptible to orogastric infection, indicating an important function for this cytokine in mucosal immunity to *T. cruzi*. To investigate the specific role of Th17 cells for mucosal immunity, we reconstituted RAG1 KO mice with *T. cruzi-specific* T cell receptor transgenic Th17 cells prior to orogastric *T. cruzi* challenges. We found that Th17 cells provided protection against gastric mucosal *T. cruzi* infection, indicated by significantly lower stomach parasite burdens. *In vitro* macrophage infection assays revealed that protection by Th17 cells is reversed with IL-17A neutralization or loss of macrophage NADPH oxidase activity. Consistent with this, *in vivo*, mice lacking functional NADPH oxidase were not protected by Th17 cell transfer. These data are the first report that Th17 cells protect against mucosal *T. cruzi* infection, and identify a novel protective mechanism involving the induction of NADPH oxidase activity in macrophages by IL-17A. These studies provide important insights for Chagas vaccine development, and more broadly, increase our understanding of the diverse roles of Th17 cells in host defense.

## Introduction

Chronic infection with the intracellular protozoan parasite *Trypanosoma cruzi* causes Chagas disease, a neglected tropical disease characterized by life-threatening cardiac and gastrointestinal pathology(1). The disease is endemic in Latin America with geographical spread into non-endemic areas, and it affects at least 8 million people(2). Infected reduviid “kissing bug” insects carry *T. cruzi* in their gastrointestinal tracts and deposit infectious parasites in their excreta after taking a blood meal. People become infected when parasite-containing excreta are accidentally ingested or inoculated into the eye or a break in the skin. Thus, the major routes of vector-borne transmission are mucosal and cutaneous.

During chronic *T. cruzi* infection, the parasite load is controlled but never completely eliminated. A robust T cell response is sufficient for parasite control in some models(3, 4), and lack of either CD4+ or CD8+ T cell responses increases susceptibility to infection(5-9). Among the CD4+ T cell subsets, Th1 cells have been demonstrated to protect against both systemic and mucosal infection(5, 10-12), while Th2 cells promote parasite persistence and mortality(12, 13). More recent studies have investigated the role of Th17 cells and IL-17A. We previously discovered using an adoptive cell transfer model that Th17 cells significantly reduce parasitemia and prevent mortality after a normally lethal challenge administered via subcutaneous injection of parasites (3). Other investigators have shown that mice deficient in IL-17A signaling due to genetic mutation of IL-17A (14) or its receptor(15), or through antibody neutralization(16), have increased susceptibility to an intraperitoneally administered *T. cruzi* challenge.

Despite evidence that IL-17A and Th17 cells protect against systemic parasite challenges, whether this type of response contributes to mucosal immunity is unknown. Orogastric infection has caused hundreds of outbreaks in humans(17-19). It has grown more common in recent years and is a leading route of transmission in some endemic areas(19). Oral ingestion is also considered to be a major route of transmission in other mammalian hosts such as domestic dogs, which are infected at high rates in endemic areas(20). Th17 cells are highly abundant at mucosal surfaces, especially in the intestinal gut, where they play a major role in homeostasis and immunity(21). Mainly through the secretion of IL-17A, these cells can recruit neutrophils, induce the expression of anti-microbial peptides, and upregulate factors that maintain epithelial integrity. These functions contribute to immunity in the intestinal mucosa(21) and may also be relevant for immunity in the gastric mucosa.

In this project, we investigated whether Th17 cells, previously demonstrated to confer immunity against systemic *T. cruzi* infection, can also drive immunity protective against mucosal infection. Ultimately, understanding of the full spectrum of functions of Th17 cells and IL-17A in *T. cruzi* infection will guide the development of vaccines inducing protective mucosal and systemic immunity.

## Results

### IL-17A signaling is important for mucosal immunity to *T. cruzi* infection

IL-17A is critical in systemic immunity against *T. cruzi* infection(14-16). To investigate how IL-17A functions in *T. cruzi* mucosal immunity, we performed orogastric infections in wild-type (WT) and IL-17RA knock-out (KO) BALB/c mice, which lack a subunit of the heterodimeric receptor for IL-17A(21). Twelve days later, we performed hematoxylin and eosin staining of tissue sections taken where *T. cruzi* preferentially infects at the margo plicatus(22), the region of transition between the glandular corpus of the stomach and the non-glandular forestomach. At baseline, the stomachs of WT and IL-17RA KO mice were histologically similar, with the presence of well-differentiated glands (Supplemental Fig 1). After infection, only mild histological abnormalities were observed in WT mice (Fig 1A). However, infected IL-17RA KO mice exhibited pronounced inflammatory infiltrate and loss of specialized glandular cells representing atrophy(23), indicating increased susceptibility to gastric mucosal parasite challenge (Figs 1A and 1B). IL-17RA KO mice also had higher parasite burdens in the stomach and a greater proportion of infected spleen cells (Figs 1C and 1D), indicating comparably worse immune control. These data are the first evidence that IL-17 signaling plays a critical role in protective gastric mucosal immunity to *T. cruzi* infection.

**Fig 1.**
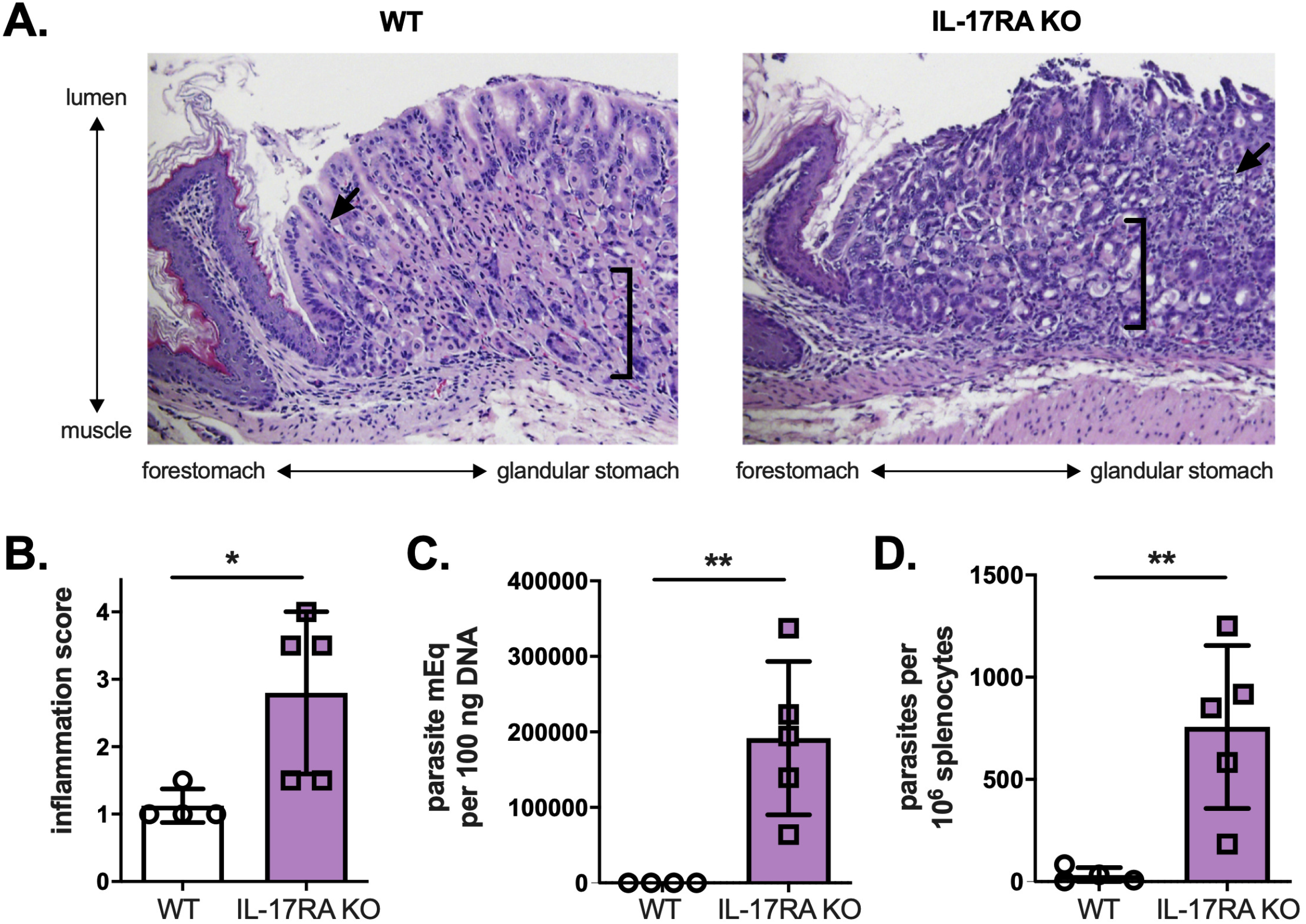
IL-17A signaling is required for mucosal immunity against gastric *T. cruzi* infection. WT and IL-17RA KO mice were orogastrically infected with *T. cruzi* parasites, then sacrificed for tissue studies 12 days later. (A) Hematoxylin and eosin staining of stomach tissue sections taken near the margo plicatus reveals greater histological disturbances and inflammation in IL-17RA KO mice. Arrows indicate areas of immune cell infiltrate, and brackets capture areas with loss of specialized cells (e.g. chief cells) at the bases of the glands (B) IL-17RA KO mice had higher inflammation scores in the gastric mucosa, based on the extent of mononuclear cell infiltrate. (C-D) IL-17RA KO mice had higher parasite DNA levels in stomach as measured by qPCR (C), and greater frequencies of infected splenocytes (D), reflecting decreased control of infection. **p<0.01, *p<0.05 by two-tailed Student t test compared to WT mice.

### Th17 cells can protect against gastric mucosal infection

Th17 cells are major CD4+ T cell producers of IL-17A. To investigate the role of Th17 cells in gastric mucosal *T. cruzi* infection, we generated Th17 cells specific for *T. cruzi*. This was done via *in vitro* Th17 differentiation of CD4+ T cells with transgenic T cell receptors (TCR Tg) recognizing an immunodominant epitope of the *T. cruzi trans-sialidase* antigen(3). We have previously characterized the phenotype and persistence of these TCR Tg Th17 cells in adoptive transfer models(3), and cells were confirmed to express the canonical Th17 cell markers RORγt and IL-17A prior to every transfer.

We transferred these TCR Tg Th17 cells into RAG1 KO mice lacking endogenous T cells, with or without *T. cruzi-naïve* polyclonal CD8+ T cells. Control animals received CD8+ T cells alone or no T cell transfer. The following day, we induced stomach infection via oral gavage of parasites. On day 12 post-infection, previously identified as the time of peak parasite burden, we sacrificed the mice for quantification of parasite burden in the stomach using qPCR and in the spleen using parasite outgrowth assays (Fig 2A). Mice receiving Th17 cells with CD8+ T cells had significantly reduced *T. cruzi* DNA in the stomach compared to control mice (Fig 2B). In addition, these mice had a smaller proportion of infected spleen cells compared to control mice, indicating significantly improved overall control of the infection (Fig 2C).

**Fig 2.**
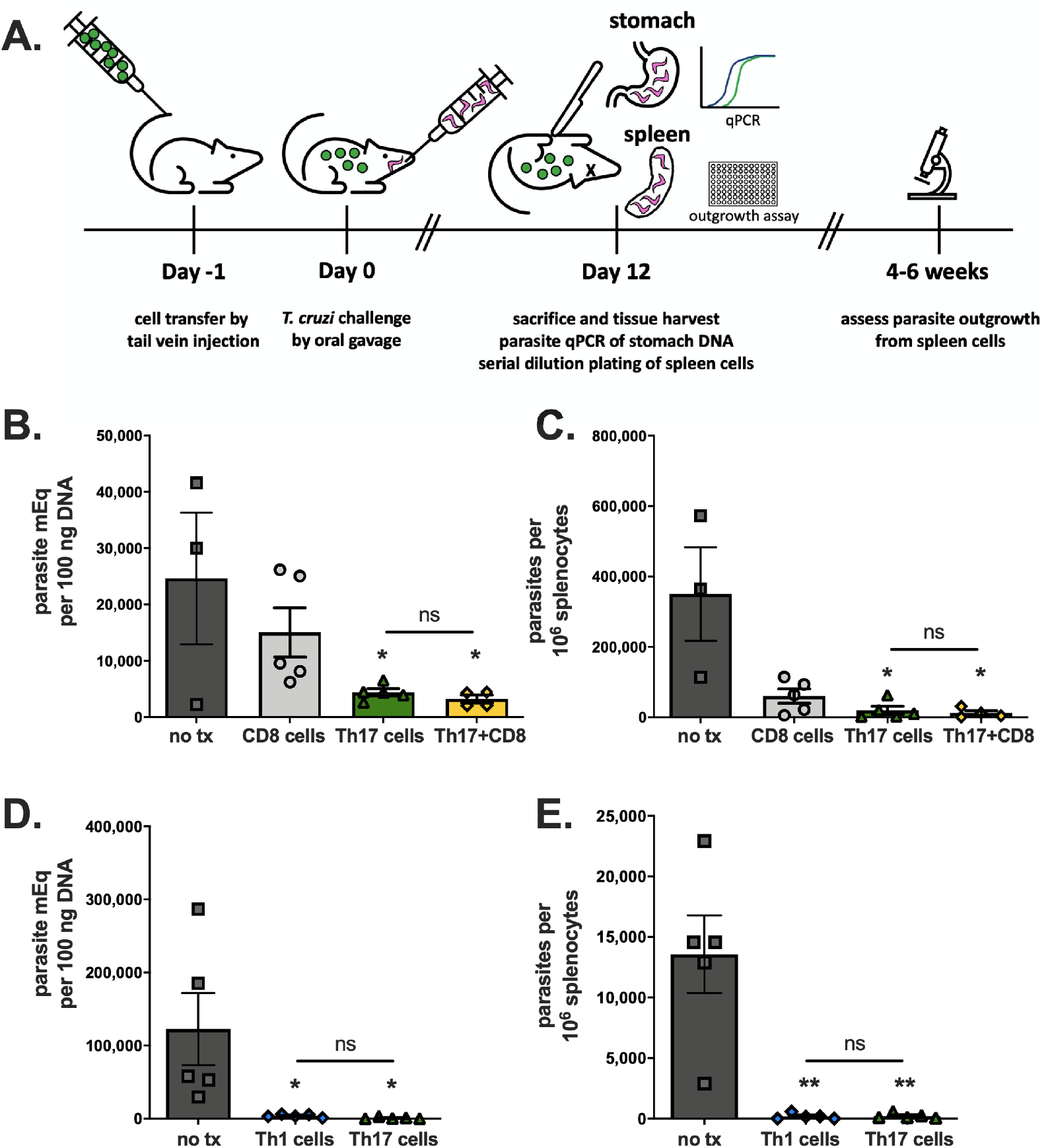
Th17 cells can provide direct protection against *T. cruzi* gastric mucosal infection. (A) RAG1 KO BALB/c mice were given various combinations of cells by adoptive transfer, orogastrically infected with *T. cruzi* parasites the following day, then sacrificed 12 days post-infection for studies of tissue parasite burden. (B-C) Mice given parasite-specific Th17 cells before challenge had decreased parasite DNA burdens in the stomach measured by qPCR (B) and a smaller frequency of infected splenocytes measured by limiting dilution parasite outgrowth assay (C). The addition of CD8+ T cells did not significantly improve this Th17-mediated protection (B-C). (D) Both Th1 and Th17 cells significantly lowered stomach parasite burdens arising after mucosal infection. (E) Th1 cells and Th17 cells comparably decreased the burden of infection in the spleen. **p<0.01, *p<0.05 by two-tailed Student t test compared to CD8 only group (B) or no tx group (C-E). Results are representative of two separate experiments.

We previously demonstrated that in systemic infection, co-transfer of CD8+ T cells is required for the protective effects of Th17 cells. Mechanistically, parasite-specific Th17 cells protected by providing help to CD8+ T cells via IL-21 signaling(3). Surprisingly, Th17 cells alone provided significant mucosal protection, and the addition of CD8+ T cells did not improve protection (Figs 2B and 2C). These data indicate that Th17-mediated mucosal protection operates via a different mechanism, involving direct protective functions rather than helper effects on CD8+ T cells.

Th1 cells are known to provide direct protective effects that can control *T. cruzi* infection independent of CD8+ T cells, primarily through IFN-γ-mediated activation of macrophages(24). We next reconstituted RAG1 KO mice with either TCR Tg Th1 or Th17 cells to compare the direct protective effects of these CD4+ T cell subsets. Both Th1 and Th17 cells were able to confer direct protective effects against a gastric challenge, as indicated by low levels of parasite DNA in the stomach after infection (Fig 2D). Th1 and Th17 cells similarly reduced the proportion of infected spleen cells compared to control mice given no T cell transfer (Fig 2E), indicating that both Th1 and Th17 cells protect against a mucosal *T. cruzi* infection even in a CD8+ T cell and B cell deficient environment.

### Th17 cells provide direct protective effects *in vitro* via IL-17A

To investigate direct protective effects of Th17 cells, we infected macrophages with *T. cruzi in vitro*, and then co-cultured them with either parasite-specific Th1 or Th17 cells before enumerating the number of infected macrophages arising after 2 days (Fig 3A). Both Th1 or Th17 cells resulted in a significantly reduced number of infected cells (Fig 3B). Adding an anti-IL-17A neutralizing antibody partially reversed the protective effects of Th17 cells (Fig 3C), suggesting that the protection is cytokine-mediated. Consistent with this, treatment with IFN-γ or IL-17A, representing the major cytokines produced by Th1 and Th17 cells, respectively, also significantly reduced the number of infected cells in both murine (Fig 3D) and human macrophages (Fig 3E). These data confirm that Th17 cells provide direct protective effects via IL-17A and suggest a similar effect may exist in humans.

**Fig 3.**
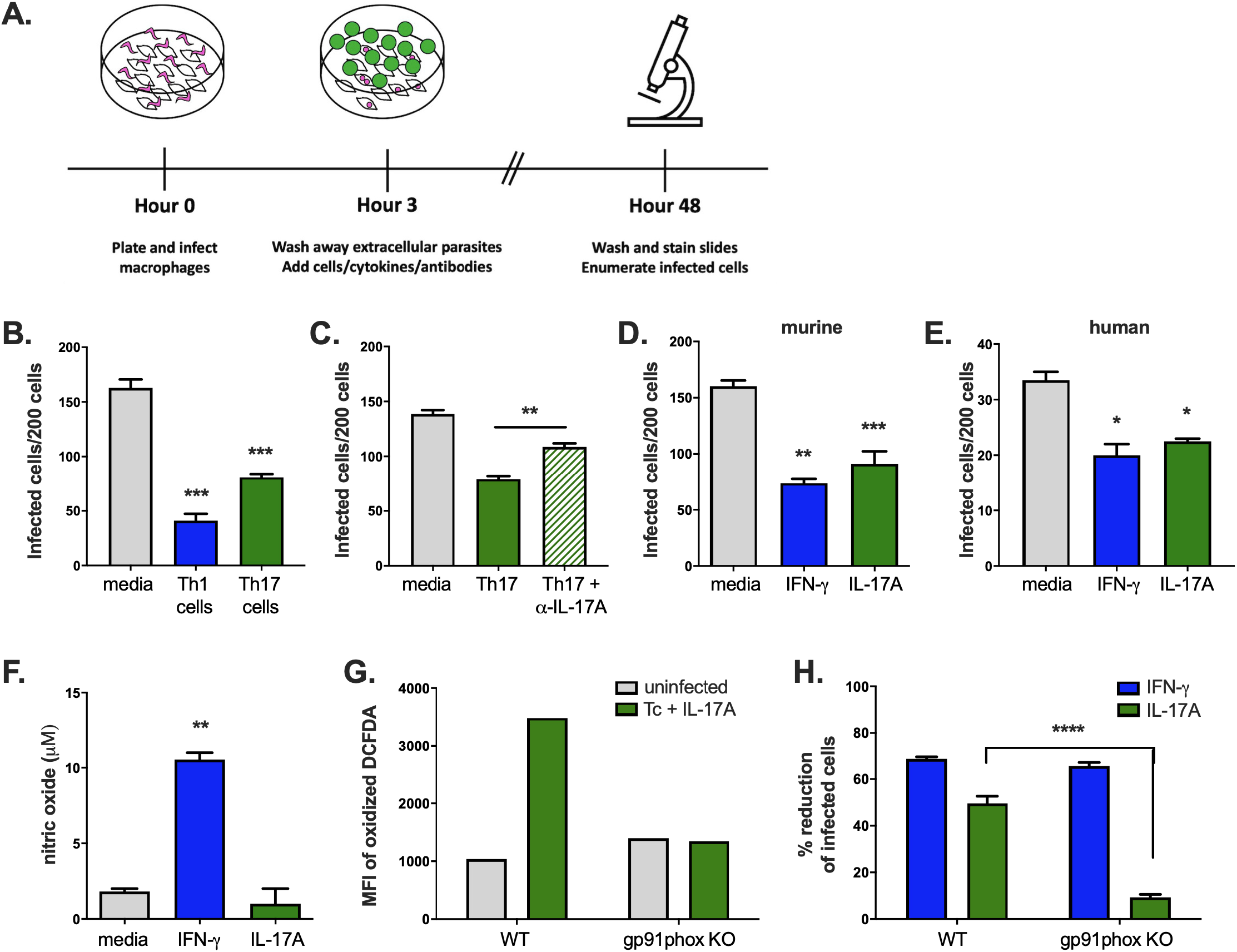
Th17 cells and IL-17A alone can inhibit *T. cruzi* intracellular growth in macrophages *in vitro* by inducing NADPH oxidase activity. (A) Macrophages were infected *in vitro* with *T. cruzi* parasites and co-cultured with T cells or cytokines. The number of infected macrophages was counted after 2 days. (B) The addition of either parasite-specific Th1 or Th17 cells reduced the number of infected cells. (C) The protective action of Th17 cells was reduced in the presence of an anti-IL-17A neutralizing antibody, indicating a cytokine-mediated effect. (D-E) IFN-γ or IL-17A treatment also decreased the number of infected cells among both murine (D) and human (E) macrophages. (F) Treatment of infected macrophages with IFN-γ, but not IL-17A, induced expression of nitric oxide. (G) The mean fluorescence intensity (MFI) of oxidized DCFDA, a measure of intracellular ROS, increased approximately 3.5-fold with *T. cruzi* infection and IL-17A treatment in WT macrophages, but not gp91phox KO macrophages. (H) IFN-γ treatment could protect both WT and gp91phox KO macrophages, while IL-17A was only able to significantly reduce the number of infected cells among WT macrophages, indicating that functional NADPH oxidase is required for IL-17A-mediated protection. Experiments in panels B-F were performed using BALB/c mice; those in panels G-H were performed in C57BL/6 background mice. ****p<0.0001, ***p<0.001, **p<0.01, *p<0.05 by two-tailed Student t test compared to medium alone. Results are representative of multiple separate experiments.

### IL-17A induces NADPH oxidase activity in infected macrophages

Th1 cells prime macrophage activation for the killing of intracellular microorganisms through IFN-γ-mediated induction of iNOS, which results in the generation of microbicidal nitric oxide (NO)(10, 24). We confirmed that treatment of infected BMDMs with IFN-γ resulted in an increase in NO concentration (Fig 3F). In contrast, treatment with IL-17A had no effect on increasing NO levels over control cells (Fig 3F). Reactive oxygen species (ROS) are generated by NADPH oxidase during the phagocyte respiratory burst response, and similar to NO, they can inhibit microbial growth. To evaluate whether ROS were induced by IL-17A, we incubated IL-17A-treated and infected BMDMs with DCFDA, a probe that can be oxidized into a fluorescent substrate. *T. cruzi* infection in the presence of IL-17A induced a 3.5-fold increase in the mean fluorescence intensity of oxidized DCFDA over uninfected cells, indicating a higher oxidation state under these conditions (Fig 3G). However, the amount of oxidized DCFDA did not increase among gp91phox KO cells lacking functional NADPH oxidase enzyme under the same conditions, reflecting the expected defect in ROS production (Fig 3G). Further, this deficiency of functional NADPH oxidase had no effect on IFN-γ-mediated protection, but reversed IL-17A-mediated protection (Fig 3H). These data indicate that direct protection by Th17 cells requires IL-17A signaling and NADPH oxidase activity.

### Th17 cells induce oxidation in mucosal immune cells *in vivo*

To determine whether parasite-specific Th17 cells given by adoptive transfer could be detected in the stomach, we recovered cells from the gastric mucosa of mice given no T cells or Th17 cells after orogastric infection with *T. cruzi*. We confirmed that parasite-specific Th17 cells were present in the gastric mucosa post-infection (Fig 4A). Proportions of other immune cell subsets, including neutrophils, which can be recruited by IL-17A, were not significantly altered compared to control mice receiving no T cell transfer (Fig 4A).

**Fig 4.**
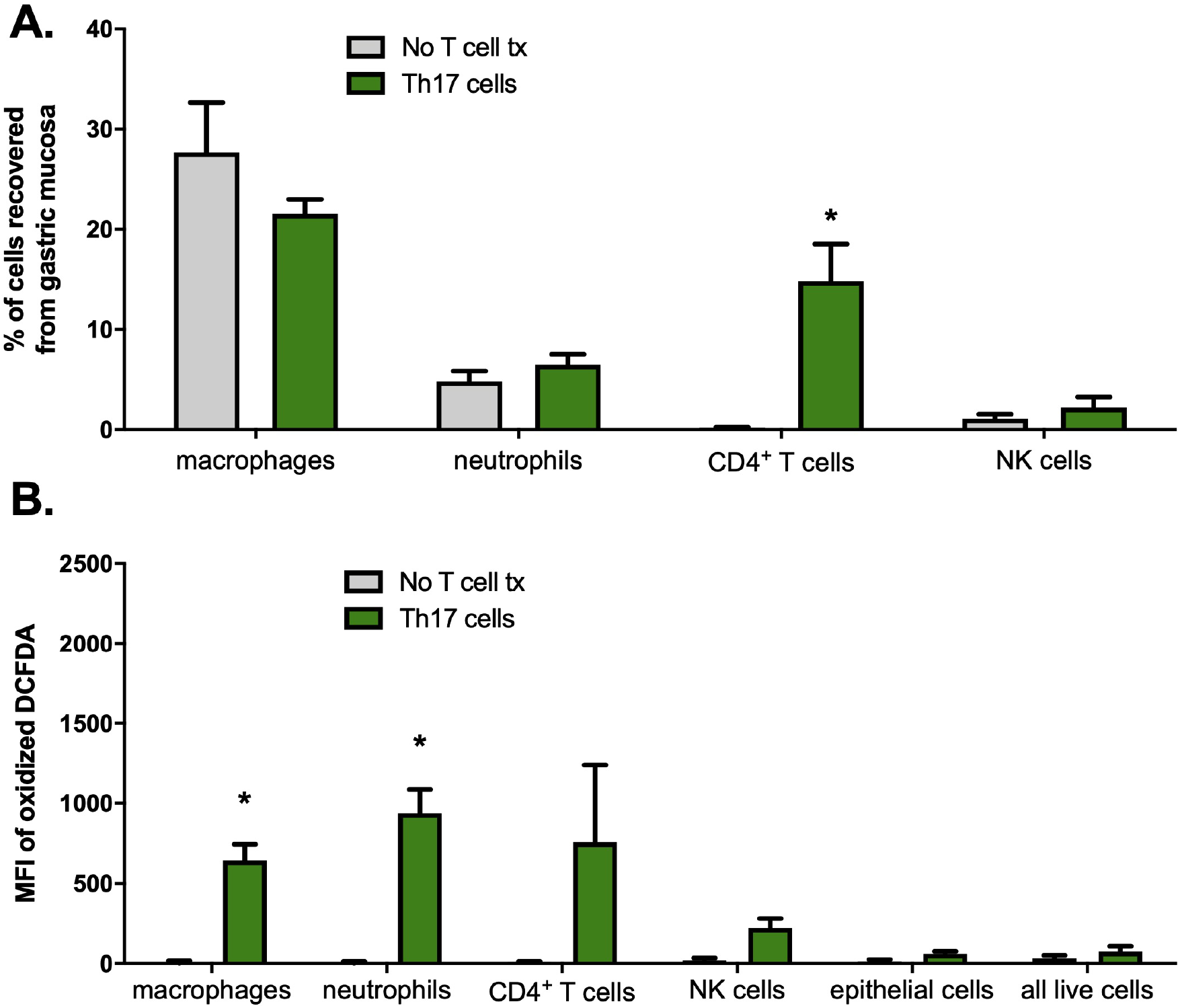
Th17 cells traffic to the gastric mucosa and induce increased expression of intracellular ROS in macrophages. RAG1 KO BALB/c mice were reconstituted with Th17 cells and orogastrically infected with *T. cruzi* the next day. Twelve days after infection, mice were sacrificed and immune cells were isolated from the gastric mucosa for flow cytometric analysis. (A) CD4+ T cells were recovered from the gastric mucosa of mice reconstituted with Th17 cells, demonstrating that these cells migrated to the site of infection. No significant differences in proportions of other immune cell subsets were detected between mice given Th17 cells or not. (B) A significantly higher oxidation state was observed in gastric mucosal macrophages and neutrophils recovered from mice given Th17 cell adoptive transfer compared to controls. n=3/group, *p<0.05 by two-tailed Student t test compared to mice given no T cell transfer. Cells were defined by the following surface markers: F4/80+ (macrophages), CD11b+Ly6G+ (neutrophils), CD3+ (CD4+ T cells), CD3-NKp46+ (NK cells), Live/Dead aqua- (all live cells), EpCAM+ (epithelial cells).

We next asked whether parasite-specific Th17 cells induced NADPH oxidase activity in the gastric mucosa during *T. cruzi* infection *in vivo*. We adoptively transferred Th17 cells into mice and, then orogastrically infected them and recovered gastric mucosal cells for DCFDA staining. We determined that macrophages and neutrophils recovered from the gastric mucosa of mice given Th17 cells had significantly higher oxidation states compared to control mice (Fig 4B), indicating an increase in reactive oxygen species. These results are consistent with phagocytic cells being the primary cell types undergoing the respiratory burst response and support the hypothesis that Th17 cells protect against orogastric infection by inducing increased expression of ROS in certain cells.

### NADPH oxidase activity is required for Th17-mediated mucosal protection

Based on the *in vitro* infection assays demonstrating protection by Th17 cells operates via secretion of IL-17A and induction of NADPH oxidase in target cells, we asked if these are also required for *in vivo* protection. We administered an anti-IL-17A neutralizing antibody every other day in Th17-reconstituted and orogastrically infected mice (Supplemental Fig 2). This partially reduced serum levels of IL-17A (Supplemental Fig 2A), but it did not reverse Th17-mediated protection (Supplemental Fig 2B), likely due to the incomplete abrogation of IL-17A activity (Supplemental Fig 2A).

To further study the role of NADPH oxidase *in vivo*, we transferred parasite-specific Th17 cells into RAG1 KO and gp91phox x RAG1 double KO (dKO) mice prior to orogastric challenge. While RAG1 KO mice had significantly improved resistance to orogastric *T. cruzi* infection following Th17 cell transfer (mean 2,783 parasite mEq per 100ng gastric DNA with Th17 cell transfer, versus 12,548 without transfer, P<0.005), Th17 cells did not confer measurable protection in mice lacking functional NADPH oxidase (mean 12,690 parasite mEq per 100ng gastric DNA with Th17 cell transfer, versus 20,252 without transfer, P=0.4304). These data indicate a critical role for this enzyme in IL-17A-mediated immunity (Fig 5).

**Fig 5.**
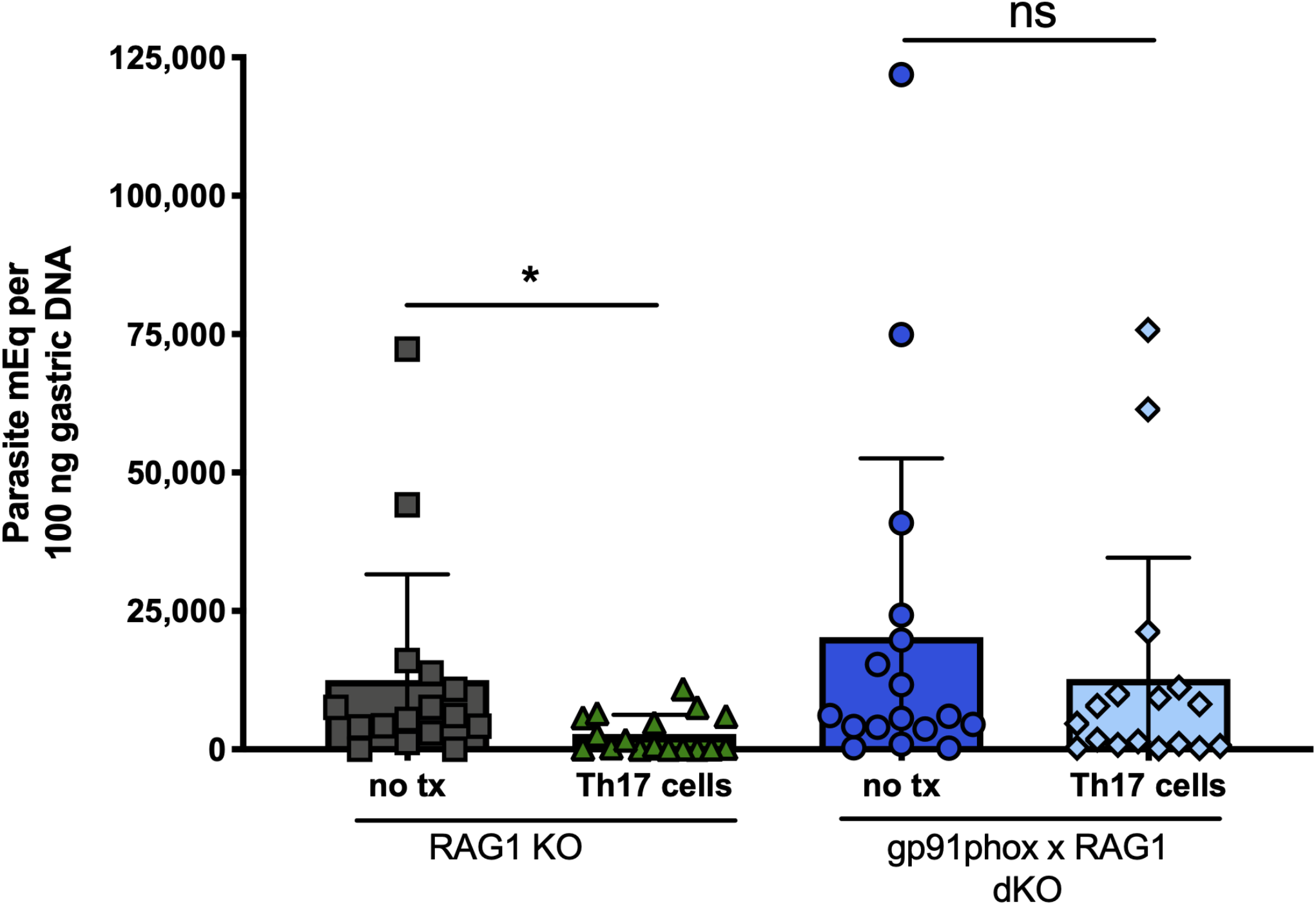
Th17 cells cannot protect in the absence of functional NADPH oxidase. RAG1 KO mice and gp91phox x RAG1 dKO C57BL/6 mice lacking functional NADPH oxidase were given parasite-specific Th17 cells or not, orogastrically challenged with *T. cruzi* the next day, then sacrificed for tissue studies 12 days after infection. Only RAG1 KO mice and not gp91phox x RAG1 dKO mice were protected from orogastric *T. cruzi* infection by parasite-specific Th17 cells, as indicated by no significant decrease in stomach parasite DNA load. *p<0.05 by two-tailed Student t test. Results depict pooled data from three independent experiments.

## Discussion

Although Chagas disease is an important illness causing significant morbidity and mortality in the Western hemisphere, few truly effective treatments exist. Drugs are limited by side effects and poor efficacy during the chronic stage of infection(25), and no vaccines are being tested in humans. The development of an effective vaccine would be an economically sound approach (26) and requires a thorough understanding of the protective immune response to this parasite, including mucosal immunity.

Th17 cells are well-established as important players against various fungi and extracellular bacteria(21) but are only recently gaining recognition as effectors in the immune response to intracellular pathogens like *T. cruzi*. Several studies over the past decade have described a role for IL-17A in the protective response against systemic *T. cruzi* parasite infection in mice(14, 16), and more recent work suggests a protective role for this cytokine in human Chagas disease as well(27). However, none of these studies has specifically examined the role of IL-17A in mucosal immunity against *T. cruzi*.

In this study, we demonstrate for the first time that Th17 cells and IL-17A contribute immunity against a gastric mucosal *T. cruzi* infection. Although CD8+ T cells are critical for Th17-mediated protection in systemic infection (3), they are dispensable in mucosal immunity. Instead, Th17 cells provide direct protective effects via secretion of IL-17A and induction of NADPH oxidase. While systemic *T. cruzi* infection involves multiple tissues and requires CD8+ T cell responses, local direct protective effects of IL-17A may be sufficient in mucosal infection. In addition, in systemic infection models, Th17 cells provided more protection against mortality compared to Th1 cells due to improved helper effects. However, protection by Th1 and Th17 cells was comparable at the gastric mucosa, where these cells act directly.

In summary, we identify a role for Th17 cells and IL-17A in mucosal immunity against *Trypanosoma cruzi* and we describe a novel protective mechanism of IL-17A against an intracellular pathogen. Th17 cells may have more broad protective functions than previously believed, and these studies in *T. cruzi* could also provide insights into infections with similar mucosally transmitted intracellular protozoa. Ultimately, these findings should be assessed for translational potential through studies targeting the induction of Th17 cells through mucosal vaccination, with the goal of reducing the significant morbidity and mortality associated with this disease.

## Materials and Methods

### Mice

*T. cruzi*-specific BALB/c mice containing a transgenic CD4+ TCR specific for p7, an immunodominant CD4 epitope of the trans-sialidase antigen, were generated in the Hoft laboratory as previously described(3). WT BALB/c mice (NCI Charles River Laboratories), RAG1 KO BALB/c mice (The Jackson Laboratory), IL-17RA KO BALB/c mice (Amgen), WT C57BL/6 (NCI Charles River), RAG1 KO C57BL/6 (The Jackson Laboratory), and B6.129S6-Cybbtm1Din/J mice lacking the gp91phox catalytic subunit of NADPH oxidase (gp91phox KO, The Jackson Laboratory) were obtained directly from the vendor or maintained as breeding colonies within the Hoft laboratory. Gp91phox KO mice were also bred to RAG1 KO C57BL/6 mice to generate RAG1 x gp91phox dKO C57BL/6 mice. All studies were approved by the Saint Louis University Institutional Animal Care and Use Committee (IACUC) under protocol #1106 and conducted in an AAALAC accredited facility at Saint Louis University. Euthanasia was performed using CO2 narcosis according to the American Veterinary Medical Association guidelines on euthanasia.

### Parasites and challenges

Tulahuèn strain parasites were maintained by *in vivo* passage through *Dipetalogaster maximus* insects and WT BALB/c mice and *in vitro* passage in LDNT+, an H_2_O-based medium containing Liver Digest Neutralized Tryptose broth (Oxford). Culture-derived metacyclic trypomastigotes (CMT) were generated by *in vitro* differentiation in supplemented Grace’s insect medium (Sigma) and maintained in a 26°C parasite incubator. For orogastric infections, mice were fasted for three hours and then fed 500 μl of a 1.5% sodium bicarbonate solution using a 22-gauge animal feeding needle, followed by 1 x 10^7^ *T. cruzi* CMT parasites in 100 μl of 1% glucose in phosphate buffered saline (PBS). Extent of infection was assessed via quantification of parasite load in the stomach and spleen. Because orogastrically infected mice do not typically mount very high blood parasite burdens compared to systemically infected mice or die due to orogastric infection, these parameters were not routinely measured in our study.

### Parasite DNA qPCR

Twelve days after orogastric infection, stomachs were dissected, cut along the greater curvature, rinsed in PBS to remove food contents, and minced with scissors. DNA was extracted from stomach tissue using DNeasy Blood and Tissue kits according to the manufacturer’s instructions (Qiagen). Primers and TaqMan probes specific for *T. cruzi* genomic DNA were used to measure parasite load in the stomach as previously described (28). When histological analyses were performed in the same experiment, the stomachs were cut along both curvatures and dorsal portions used for qPCR.

### Histology

Longitudinal strips 2-3 mm in width were cut from the esophageal to the pyloric ends of the ventral stomach, placed between pre-wetted foam pads in biopsy cassettes (Leica), and fixed by submersion in 10% neutral buffered formalin. Several hours later, the tissue was embedded, sectioned, affixed to slides and stained with hematoxylin and eosin at the Saint Louis University Research and Histology Microscopy Core. Images were acquired on an Olympus BX41 or Leica epifluorescent microscope. Inflammation scores were assigned according to a qualitative 0-4 scale by a blinded evaluator based on a scoring system adapted from Rogers et al(29). Briefly, inflammation was scored based upon the presence of leukocytes and the extent of their infiltration into the gastric mucosa. A score of 0 would indicate only the number of leukocytes present in the tissue at baseline (i.e. in a normal uninfected mouse with no inflammation), while a score of 4 indicates severe transmural inflammation with the presence of numerous infiltrating leukocytes.

### Generation of Th1 and Th17 cells

*T. cruzi*-specific Th1 and Th17 cells were generated *in vitro* as previously described (3). Briefly, TCR Tg CD4+ T cells were stimulated with irradiated, cognate peptide-pulsed dendritic cells in the presence of α-IL-4 and IL-12 (to generate Th1 cells), or α-IL-4, α-IFN-γ, IL-6, TGF-β, and IL-23 (to generate Th17 cells). Media was refreshed on days 3 and 6 with IL-2 for Th1 cells or IL-23 for Th17 cells. The cells underwent two rounds of this differentiation (re-stimulation with peptide-pulsed APCs on day 7 followed by maintenance cytokines on days 10 and 13). Differentiation was confirmed by intracellular cytokine staining (ICS) for transcription factors and cytokines, and the cells were used on day 14. The phenotypes and stability of these Th1 and Th17 cells is previously documented(3).

### Flow cytometry and intracellular cytokine staining (ICS)

To obtain gastric mucosal immune cells, stomachs were dissected open, a syringe needle was introduced into the mucosal layers, and the region flushed with large volumes of media. The recovered cells were cultured with 1 μl/ml of GolgiPlug containing monensin (BD Pharmingen) and 0.67 μl/ml of GolgiStop containing Brefeldin A (BD Pharmingen) for 3 hours at 37°C. Cells were then stained with Live Dead fixable aqua, followed by surface antibodies directed against CD3, CD4, CD11b, CD11c, Ly6G, F4/80, NKp46 (all BD or eBiosciences). Cells were fixed and permeabilized for intracellular cytokine staining using a Foxp3 transcription buffer staining set (eBiosciences). Cells were washed with 1% FBS in Dulbecco’s PBS between all steps. All incubations were performed for 30 minutes at 4°C. For oxidation staining, cells were incubated for 20 minutes at 37°C with 100 nM of H_2_DCFDA (Invitrogen) after surface staining. All samples were acquired on a BD LSRII flow cytometer at the Saint Louis University flow cytometry core facility and analyzed on FlowJo software (Treestar, Inc.).

### Splenic parasite outgrowth assay

Total spleen cells were mechanically isolated from dissected spleens 12 days after orogastric infection. Spleen cells were re-suspended in LDNT+ parasite medium, plated in a limiting dilution fashion in 96-well plates, and incubated in a 26°C parasite incubator. One month later, all wells were examined for live parasites. The last well with parasite outgrowth was used to estimate the number of parasites per million cells (e.g., live parasites emerging from 100 plated cells = at least 1 parasite per 100 cells) (28). Because a single cell suspension can be obtained from the spleen, this method was selected over qPCR for its ability to quantify the presence of live parasites, though it may underestimate the true burden due to the detection of only parasites that can differentiate into trypomastigotes.

### Generation of peritoneal exudate macrophages (PEMs), bone-marrow derived macrophages (BMDMs), and human monocyte-derived macrophages

PEMs were generated from BALB/c strain mice, and BMDMs were generated from C57BL/6 strain mice. For PEMs, mice were injected intraperitoneally with 100 μg of concanavalin A (Sigma). PEMs were recovered 3-4 days later via peritoneal lavage using a syringe and needle. To generate BMDM, bone marrow cells harvested from mouse femurs were plated in supplemented RPMI medium with 20 ng/ml M-CSF (eBioscience), refreshed on day 3. BMDMs were harvested for use on day 7. For human monocyte-derived macrophages, peripheral blood mononuclear cells were cultured in 96-well plates for 7 days to induce maturation. Adherent cells were removed and re-plated in 8-well chamber slides (Lab-Tek) for infection assays.

### *In vitro* infection assay

Macrophages were plated at 200,000 cells/well of 8-well chamber slides and infected with CMT (MOI=10). After 3 hours, extracellular parasites were removed by repeated washing. In some experiments, macrophages were co-cultured with purified recombinant murine or human IFN-γ (Genentech) at 1000 U/ml or IL-17A (R&D) at 100 ng/ml, or an anti-IL-17A neutralizing antibody (TC11-18H10, BD Pharmingen) was added at a concentration of 10 μg/ml. Nitric oxide concentrations in supernatants were measured using the Griess Reagent System (Promega). Slides were stained with Diff-Quik (IMEB, Inc.) after 2 days, and infected cells were enumerated microscopically.

### *In vivo* IL-17A neutralization

Mice were given 100 μg of an anti-IL-17A neutralizing antibody (17F3, Bio X Cell) or an IgG1 isotype control antibody (MOPC-21, Bio X Cell) every 2 days by intraperitoneal injection. Serum IL-17A levels were analyzed via Mouse IL-17A ELISA MAX Standard kits (BD).

## Statistics

All analyses were performed in Prism version 8 (GraphPad Software) using a significance level of 5%.

## Acknowledgements

The authors thank Jennifer Franey for her assistance with animal care and handling; Joy Eslick and Sherri Koehm for their assistance with flow cytometry; and Grant Kolar, Barbara Nagel and Caroline Murphy for their assistance with histology and microscopy. We thank Amgen for the provision of IL-17RA KO mice received under a material transfer agreement.

## Author contributions

**Conceptualization:** CWC, CSE, DFH

**Formal analysis:** CWC, CSE, KAM, JRB, REA, DHC, KAB, DFH

**Funding acquisition:** DFH, CWC

**Investigation:** CWC, CSE, KAM, JRB, REA, DHC, KAB

**Methodology:** CWC, CSE, JRB, KAB, RJD, DFH

**Supervision:** DFH

**Validation:** CWC, CSE, KAM, JRB, REA, DHC

**Visualization:** CWC

**Writing – original draft:** CWC

**Writing – revising & editing:** all authors

---

**Supplemental Fig 1.**
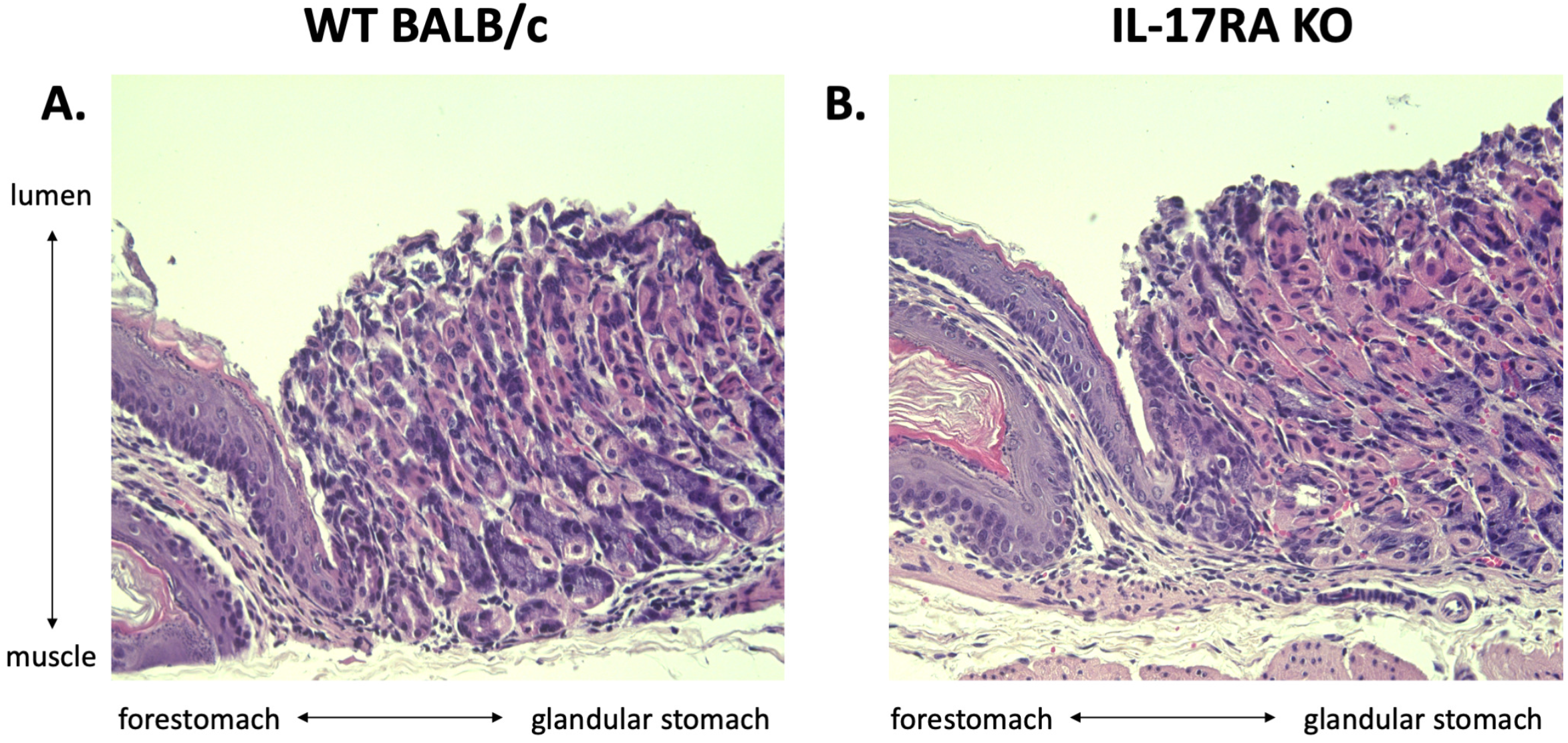
Stomachs of WT and IL-17RA KO BALB/c mice are similar at baseline. Prior to infection, both WT (A) and IL-17RA KO (B) mice demonstrated the presence of well-differentiated gastric glands and no significant immune cell infiltrate in the stomach mucosa.

**Supplemental Fig 2.**
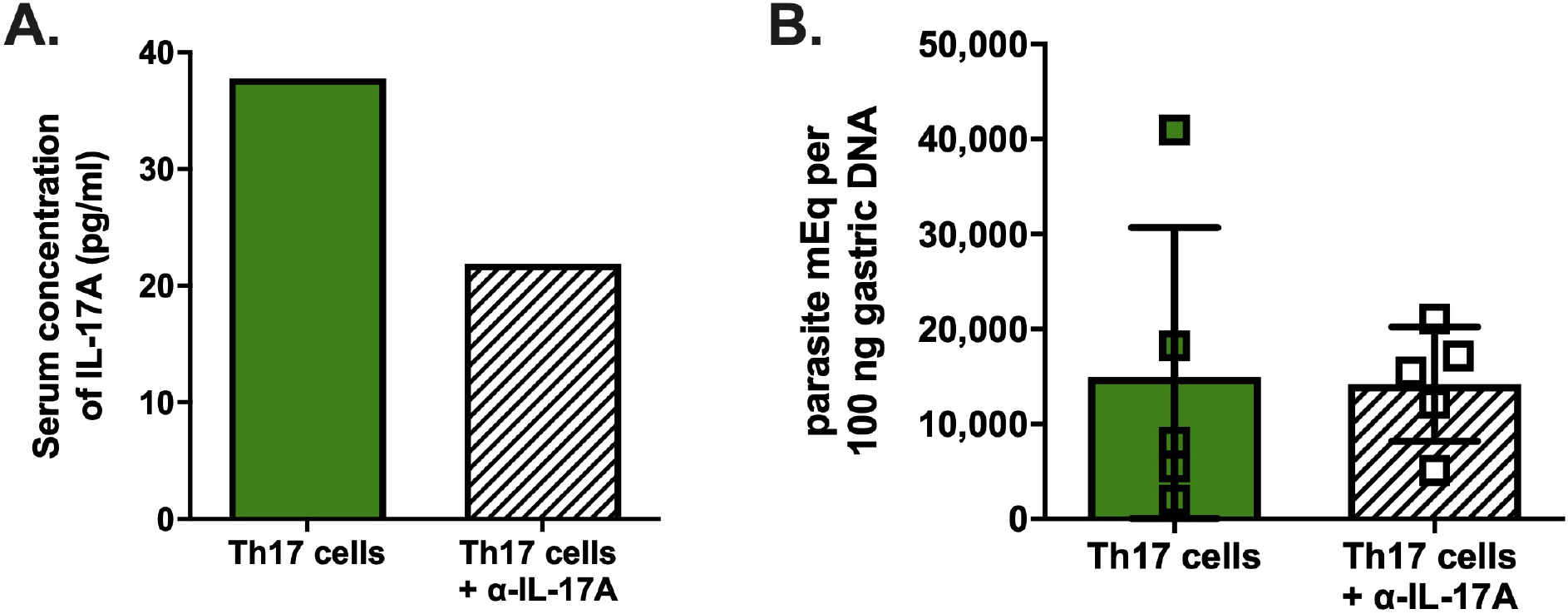
IL-17A neutralization does not reverse protection by Th17 cells *in vivo*. RAG1 KO BALB/c mice reconstituted with parasite-specific Th17 cells were orogastrically infected with *T. cruzi* the next day and treated with an anti-IL-17A neutralizing antibody or an isotype control antibody every other day until sacrifice on day 12 post-infection for tissue studies. (A) Pooled sera of mice treated with anti-IL-17A show reduced but not abolished serum IL-17A. (B) Treatment with anti-IL-17A had no effect on Th17-mediated suppression of gastric parasite burden after orogastric infection.

## References

1. Rassi A, Jr., Rassi A, Marin-Neto JA. Chagas disease. Lancet. 2010;375(9723):1388–402.

2. Bern C, Montgomery SP. An estimate of the burden of Chagas disease in the United States. Clin Infect Dis. 2009;49(5):e52–e4.

3. Cai CW, Blase JR, Zhang X, Eickhoff CS, Hoft DF. Th17 Cells Are More Protective Than Th1 Cells Against the Intracellular Parasite Trypanosoma cruzi. PLoS Pathog. 2016;12(10):e1005902.

4. Sullivan NL, Eickhoff CS, Sagartz J, Hoft DF. Deficiency of Antigen-Specific B Cells Results in Decreased *Trypanosoma cruzi* Systemic but Not Mucosal Immunity Due to CD8 T Cell Exhaustion. J Immunol. 2015;15(194):1806–18.

5. Hoft DF, Eickhoff CS. Type 1 immunity provides both optimal mucosal and systemic protection against a mucosally invasive, intracellular pathogen. Infect Immun. 2005;73(8):4934–40.

6. Tarleton RL, Sun J, Zhang L, Postan M. Depletion of T-cell subpopulations results in exacerbation of myocarditis and parasitism in experimental Chagas’ disease. Infect Immun. 1994;62(5):1820–9.

7. Tarleton RL, Koller BH, Latour A, Postan M. Susceptibility of beta 2-microglobulin-deficient mice to *Trypanosoma cruzi* infection. Nature. 1992;356(6367):338–40.

8. Tarleton RL, Grusby MJ, Postan M, Glimcher LH. Trypanosoma cruzi infection in MHC-deficient mice: further evidence for the role of both class I- and class II-restricted T cells in immune resistance and disease. Int Immunol. 1996;8(1):13–22.

9. Tarleton RL. Depletion of CD8^+^ T cells increases susceptibility and reverses vaccine-induced immunity in mice infected with *Trypanosoma cruzi*. J Immunol. 1990;144(2):717–24.

10. Hoft DF, Schnapp AR, Eickhoff CS, Roodman ST. Involvement of CD4(+) Th1 cells in systemic immunity protective against primary and secondary challenges with *Trypanosoma cruzi*. Infect Immun. 2000;68(1):197–204.

11. Hoft DF, Eickhoff CS. Type 1 immunity provides optimal protection against both mucosal and systemic *Trypanosoma cruzi* challenges. Infect Immun. 2002;70(12):6715–25.

12. Kumar S, Tarleton RL. Antigen-specific Th1 but not Th2 cells provide protection from lethal *Trypanosoma cruzi* infection in mice. J Immunol. 2001;166(7):4596–603.

13. Lopes MF, Nunes MP, Henriques-Pons A, Giese N, Morse HC, 3rd, Davidson WF, et al. Increased susceptibility of Fas ligand-deficient gld mice to Trypanosoma cruzi infection due to a Th2-biased host immune response. Eur J Immunol. 1999;29(1):81–9.

14. Miyazaki Y, Hamano S, Wang S, Shimanoe Y, Iwakura Y, Yoshida H. IL-17 is necessary for host protection against acute-phase Trypanosoma cruzi infection. J Immunol. 2010;185(2):1150–7.

15. Tosello Boari J, Amezcua Vesely MC, Bermejo DA, Ramello MC, Montes CL, Cejas H, et al. IL-17RA signaling reduces inflammation and mortality during Trypanosoma cruzi infection by recruiting suppressive IL-10-producing neutrophils. PLoS Pathog. 2012;8(4):e1002658.

16. da Matta Guedes PM, Gutierrez FR, Maia FL, Milanezi CM, Silva GK, Pavanelli WR, et al. IL-17 produced during Trypanosoma cruzi infection plays a central role in regulating parasite-induced myocarditis. PLoS Negl Trop Dis. 2010;4(2):e604.

17. Nobrega AA, Garcia MH, Tatto E, Obara MT, Costa E, Sobel J, et al. Oral transmission of Chagas disease by consumption of acai palm fruit, Brazil. Emerg Infect Dis. 2009;15(4):653–5.

18. Alarcon de Noya B, Diaz-Bello Z, Colmenares C, Ruiz-Guevara R, Mauriello L, Zavala-Jaspe R, et al. Large urban outbreak of orally acquired acute Chagas disease at a school in Caracas, Venezuela. J Infect Dis. 2010;201(9):1308–15.

19. Shikanai-Yasuda MA, Carvalho NB. Oral transmission of Chagas disease. Clin Infect Dis. 2012;54(6):845–52.

20. Montenegro VM, Jimenez M, Dias JC, Zeledon R. Chagas disease in dogs from endemic areas of Costa Rica. Mem Inst Oswaldo Cruz. 2002;97(4):491–4.

21. Korn T, Bettelli E, Oukka M, Kuchroo VK. IL-17 and Th17 Cells. Annu Rev Immunol. 2009;27:485–517.

22. Hoft DF, Farrar PL, Kratz-Owens K, Shaffer D. Gastric invasion by Trypanosoma cruzi and induction of protective mucosal immune responses. Infect Immun. 1996;64(9):3800–10.

23. Fox JG, Wang TC. Inflammation, atrophy, and gastric cancer. J Clin Invest. 2007;117(1):60–9.

24. Rodrigues MM, Ribeirao M, Boscardin SB. CD4 Th1 but not Th2 clones efficiently activate macrophages to eliminate Trypanosoma cruzi through a nitric oxide dependent mechanism. Immunol Lett. 2000;73(1):43–50.

25. Pinazo MJ, Munoz J, Posada E, Lopez-Chejade P, Gallego M, Ayala E, et al. Tolerance of benznidazole in treatment of Chagas’ disease in adults. Antimicrob Agents Chemother. 2010;54(11):4896–9.

26. Lee BY, Bacon KM, Connor DL, Willig AM, Bailey RR. The potential economic value of a Trypanosoma cruzi (Chagas disease) vaccine in Latin America. PLoS Negl Trop Dis. 2010;4(12):e916.

27. Magalhaes LM, Villani FN, Nunes Mdo C, Gollob KJ, Rocha MO, Dutra WO. High interleukin 17 expression is correlated with better cardiac function in human Chagas disease. J Infect Dis. 2013;207(4):661–5.

28. Hoft DF, Farrar PL, Kratz-Owens K, Shaffer D. Gastric invasion by *Trypanosoma cruzi* and induction of protective mucosal immune responses. Infect Immun. 1996;64(9):3800–10.

29. Rogers AB, Taylor NS, Whary MT, Stefanich ED, Wang TC, Fox JG. Helicobacter pylori but not high salt induces gastric intraepithelial neoplasia in B6129 mice. Cancer Res. 2005;65(23):10709–15.

